# Morphological Metrics of Magnetic Resonance Imaging of Glioblastoma as Biomarkers of Prognosis

**DOI:** 10.1101/2021.01.29.428875

**Authors:** Lee Curtin, Paula Whitmire, Haylye White, Maciej M. Mrugala, Leland S. Hu, Kristin R. Swanson

**Author notes:** **Corresponding Author Contact:** Lee Curtin, Ph.D. Co-senior authors contributed equally (Leland S. Hu, Kristin R. Swanson). **Address:** All authors - 5777 E Mayo Blvd, Phoenix, AZ, 85054.

## Abstract

Morphological characteristics have been linked to outcomes across a variety of cancers. Lacunarity is a quantitative morphological measure of how shapes fill space while fractal dimension is a morphological measure of the complexity of pixel arrangement. Glioblastoma is the most aggressive primary brain tumor with a short expected survival given the current standard-of-care treatment. Due to the sensitive location of the tumor, there is a heavy reliance on imaging to assess the state of the disease in the clinic. In this project, we computed lacunarity and fractal dimension values for glioblastoma-induced abnormalities on gadolinium-enhanced T1-weighted magnetic resonance imaging (T1Gd MRI) as well as T2-weighted (T2) and fluid-attenuated inversion recovery (FLAIR) MRIs. In our patient cohort (n=402), we aim to connect these morphological metrics calculated on pretreatment MRI with the survival of patients with GBM. We calculated lacunarity and fractal dimension across all MRI slices on necrotic regions (n=390) and abnormalities on T1Gd MRI (n=402), as well as on enhancing abnormalities present on T2/FLAIR MRI (n=257). We also explored the relationship between these metrics and age at diagnosis, as well as abnormality volume. We found statistically significant relationships to outcome across all three imaging subtypes, with the shape of T2/FLAIR abnormalities showing the strongest relationship with overall survival. The link between morphological and survival metrics could be driven by underlying biological phenomena, tumor location or microenvironmental factors that should be further explored.

## Introduction

Glioblastoma (GBM) is an aggressive and highly infiltrative primary brain tumor with a median survival of only 15-16 months with standard-of-care treatment (1–3). Due to the sensitive location of the tumor, opportunities for biopsies are limited and there is a heavy reliance on imaging, typically magnetic resonance imaging (MRI), to assess the severity and progression of the disease. There remains a relative lack of studies on the prognostic implications of the such shape metrics of GBM-induced abnormalities on MRI. There are three key GBM-associated regions detectable on standard-of-care MRI. The first is the enhancing region present on T1-weighted MRI with gadolinium contrast (T1Gd MRI), caused through a leakage of gadolinium through disrupted vasculature (4). This enhancement typically spatially correlates with the bulk of the tumor, particularly in a pretreatment setting (5). The second region is necrosis, caused by a lack of sufficient nutrients and necrosis-inducing factors, typically present as a central hypointense region on T1Gd MRI surrounded by enhancement. The third is the abnormal region present on both T2-weighted and fluid attenuated inversion recovery (FLAIR) MRIs that spatially correlates to edema and infiltrative tumor cells (6).

In this work, we focus on the prognostic impact of two morphological metrics that quantify these GBM regions. The first is fractal dimension, a measure of the consistency of a shape with itself at varying spatial scales. If a shape is extremely self-consistent, it will have a high fractal dimension. Lacunarity is a quantifiable measure of how shapes fill space, and more generally considers heterogeneity. Higher lacunarity values occur in shapes that are disconnected and more heterogeneous. There are many examples of fractal dimension and lacunarity providing clinical insight in oncology. For example, fractal dimension and lacunarity have been shown as prognostic markers for melanoma and laryngeal carcinoma (7,8). Differences in fractal dimension between healthy and pathological tissue have been found in renal chromophobe carcinoma (9). Fractal dimension has been shown to distinguish benign and malignant breast tumors in both digitized histology (10) and ultrasound images (11). There is also a wealth of morphological studies on lung cancers (12). Within glioma, the lacunarity of T2-weighted MRI abnormalities has been shown to distinguish glioma grade (13) and the fractal dimension of brain vasculature in susceptibility-weighted imaging has been shown to distinguish brain tumor grade (14,15). A separate study of 95 patients has found fractal dimension and lacunarity, applied to pretreatment necrotic regions present on T1Gd MRIs can distinguish overall survival (OS) and progression free survival (PFS) in GBM (16). We seek to build on the results of this previous study on a larger cohort through the inclusion of analyses on other MRI abnormalities. We will also look for correlations between these morphological metrics with patient age, patient sex and imaging abnormality volumes.

In a retrospective cohort of 402 patients with newly-diagnosed GBM, we have calculated lacunarity and fractal dimension values of imaging abnormalities using T1Gd MRI and T2/FLAIR images. We find statistically significant relationships between these morphological metrics applied to imaging abnormalities and survival metrics in patients with GBM, both for OS and PFS, some of which remain significant when adjusting for multiple comparisons.

## Methodology

### Patient Cohort

We queried our multi-institutional database of retrospective patient data for patients with first-diagnosed GBM. We required these patients to have available segmented pretreatment T1-weighted MRI with gadolinium contrast in our database, as well as age at diagnosis, sex and overall survival (confirmed death, alive, or lost to follow-up). We also required that these patients’ tumors did not contain a significant cystic component, which typically presents as hypointense with surrounding enhancement and smooth on T2-weighted MRI. Cystic components are typically round and may provide a survival benefit to patients (17), which may have interfered with relationships between morphological metrics and survival. Hypointense cystic fluid would also interfere with our ability to consistently capture necrotic regions, which are also hypointense. This resulted in a cohort of 402 patients. Where available, we also noted progression-free survival (n=129) and stored T2/FLAIR segmentations (n=257). As many of our patients were diagnosed before the current standard of care (SOC) protocol was established, our cohort consists of a variety of treatment protocols. We have established a subcohort of 141 patients known to have received the current SOC, and refer to these as “current SOC patients”. See **Table 1** for further breakdowns of cohort sizes across different imaging regions, and **Supplement 1** for a sex-specific breakdown of these groups.

**Table 1:**
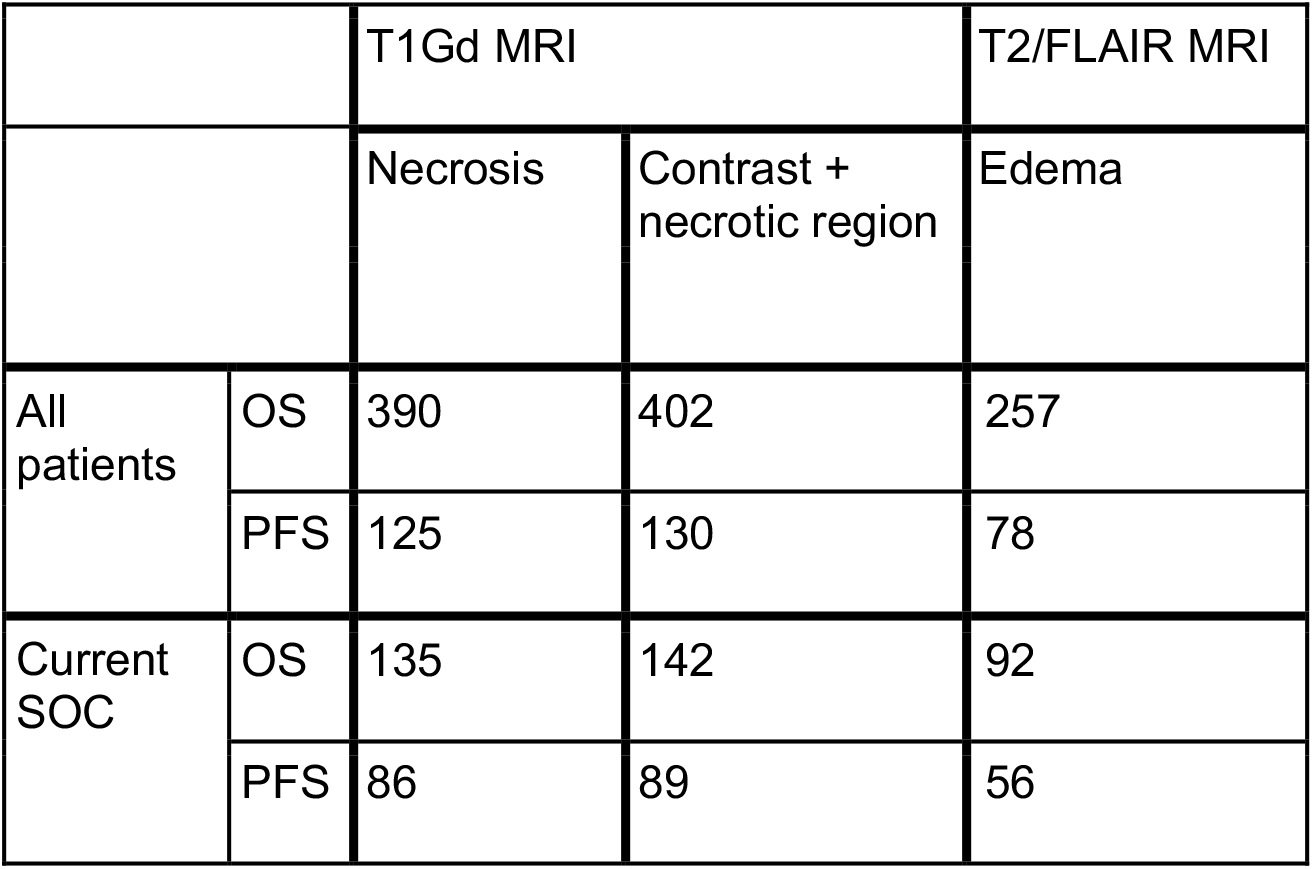
Cohort of patients with known overall survival (OS) and progression free survival (PFS). This table shows the number of patients with available imaging ROIs and known OS and PFS, including the subsets known to have received the current standard of care (SOC). The discrepancy between patients with available necrosis ROIs and T1Gd enhancing ROIs is due to 12 patients with negligible necrosis that precluded addition of those tumors to the necrosis-specific analysis.

|  |  | T1Gd MRI |  | T2/FLAIR MRI |
| --- | --- | --- | --- | --- |
|  |  | Necrosis | Contrast + necrotic region | Edema |
| All patients | OS | 390 | 402 | 257 |
|  | PFS | 125 | 130 | 78 |
| Current SOC | OS | 135 | 142 | 92 |
|  | PFS | 86 | 89 | 56 |

### Biomedical Imaging and ROI Segmentation

Using pretreatment T1Gd MRI, enhancing abnormalities were segmented by trained individuals and necrotic regions were segmented using the segmentations of T1Gd enhancement within an automated dilation erosion algorithm. T2/FLAIR abnormalities were also segmented by trained individuals. Segmentation regions were used alongside image dimensions to calculate imaging abnormality volumes (cm^3^). We also use segmented volumes to compute radii of equivalent spherical volumes (cm). There are three different segmentations used in this analysis, the first is necrotic regions, the second is T1Gd enhancing regions with necrotic regions, and the third are abnormal regions on T2/FLAIR MRI. We chose to include the necrotic regions with the enhancement on T1Gd MRI to avoid the conflation of necrosis outlines that would otherwise be present in T1Gd-enhancing regions alone.

### Lacunarity and Fractal Dimension

We used the FracLac plugin for ImageJ to calculate lacunarity and fractal dimension values for each 2D segmentation (18,19). Image slices with segmentations totalling 5 pixels or less were excluded. The FracLac software uses a box-counting algorithm to compute lacunarity and fractal dimension; we used a minimum of 2 pixels and a maximum of 45% of each MRI slice for these box sizes. To compute fractal dimension, grids with varying box sizes are placed over a region, and the number of boxes needed to cover the region in question is recorded for each grid. The log of this number is then plotted against the log reciprocal of the length of each box and the gradient of the regression line for this plot is the fractal dimension. For lacunarity, a similar process is undertaken with the numbers of pixels in each box recorded as a distribution for each box length, with standard deviation σ and mean μ. Lacunarity is then the mean over all box sizes of (σ/μ)^2. Due to the dependency of fractal dimension and lacunarity on grid placement, we used a mean fractal dimension and a mean lacunarity calculated over 12 grid placements for each 2D segmentation. We then stored the following metrics for both fractal dimension and lacunarity across MRI slices, to give one value of each per pretreatment MRI: the median value, the mean value, the range of values and the variance of values. We see the clearest survival signals for the median value, and present these in this work, with the mean of necrosis ROIs presented in **Supplement 2** to compare more closely to previously published work (16). In **Figure 1A**, we present schematics of how we compute lacunarity and fractal dimension values.

**Fig 1.**
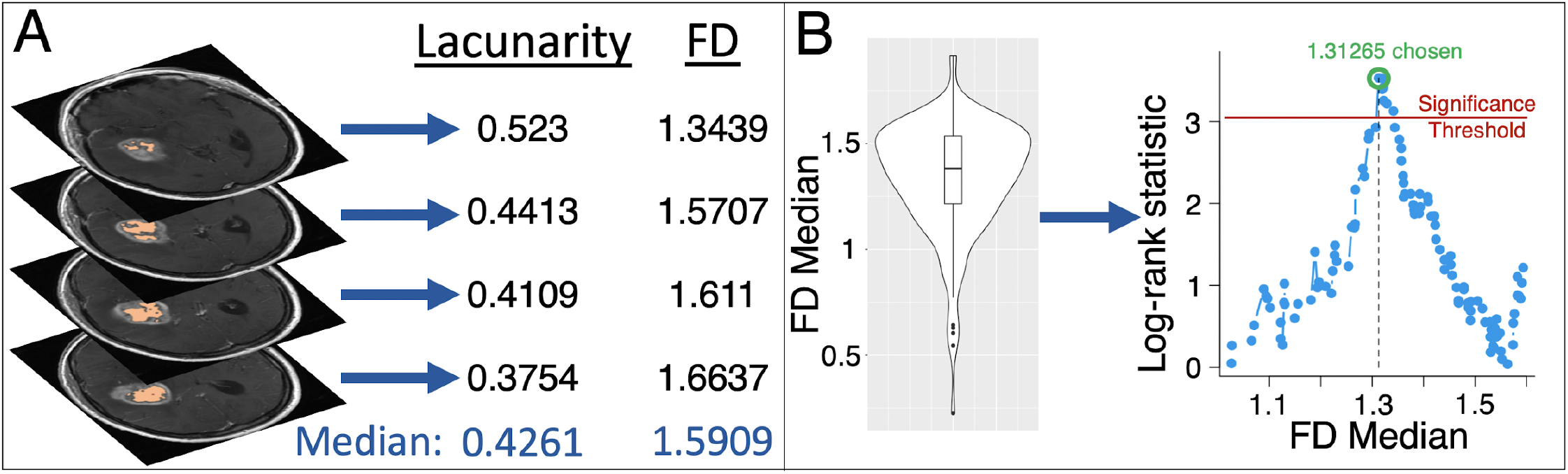
Computing lacunarity and fractal dimension (FD) and testing the statistical significance of these against patient survival data. **(A)** As an example, lacunarity and FD are computed on each slice of necrosis segmentations. The median of these values is stored, giving one value of lacunarity and FD for each patient. We do the same for T1Gd and T2/FLAIR regions. **(B)** Individual median values are collated into a cohort analysis. Each median value that does not split the cohort into groups of less than 10% of the cohort size is then tested as a cutoff to distinguish either overall survival or progression free survival. A log-rank test provides a significance value for each potential cutoff and a log-rank statistic is calculated alongside a separate significance threshold that accounts for multiple comparisons. The maximal log-rank statistic is chosen as this maximally distinguishes the two groups. This is carried out for lacunarity and FD, across necrosis, T1Gd and T2/FLAIR regions.

### Statistical Analyses

We used log-rank tests to ascertain the significance in overall survival and progression-free survival differences in our cohort; we use Kaplan-Meier curves to visualize these differences. All of the analysis presented here was carried out in R (20–24). Throughout this work, we set a p-value threshold of 0.05, below which we consider our results to be statistically significant. As this work uses multiple comparisons to look for significant thresholds that discriminate our cohorts by survival, we have used the maxstat package in RStudio to adjust our p values appropriately. Namely, we have implemented the adjustment method first presented by Lausen and Schumacher (25). We use the chosen threshold to divide our cohorts into two groups: the first consists of patients with values lower or equal to the threshold and the second consists of patients above the chosen threshold. Only thresholds that split the cohort into groups larger than 10% of the cohort size were tested. We present an example schematic of this process in **Figure 1B**. We present adjusted and unadjusted p values within this work, and will clearly state when each is used (23).

Cox proportional hazard (CPH) models have been used to test the significance of lacunarity and fractal dimension as continuous predictors of survival in univariate and multivariate analyses against values that exist within all patients such as age at diagnosis and tumor radius. We used Pearson correlation coefficient tests to determine the significance and strength of correlations between variables. We used Welch’s t-tests to determine the significance between means of different groups.

### Ethical Approval

All procedures performed in the studies involving human participants were in accordance with the ethical standards of the institutional and/or national research committee and with the 1964 Helsinki declaration and its later amendments or comparable ethical standards. Our de-identified data repository of patients with brain cancer includes retrospective data collected from medical records and prospective data collection. Research conduct on the data repository is approved by Mayo Clinic Institutional Review (17-009688). Retrospective inclusion as well informed consent was obtained for all prospectively enrolled participants in the repository as approved by Mayo Clinic Institutional Review Board (IRB# 17-009682).

## Results

### Whole Cohort

#### Necrotic Regions

We found that lacunarity significantly distinguished overall survival with the group of lower lacunarity values showing benefit but this result did not hold while adjusting for multiple comparisons (unadjusted p=0.012, adjusted p=0.07541, n=390). We also found that lacunarity significantly distinguished progression free survival. Lacunarity could still significantly distinguish PFS while accounting for multiple comparisons (adjusted p=0.0051, n=125), with higher values showing a survival benefit; this result is presented in **Figure 2A**.

**Fig 2.**
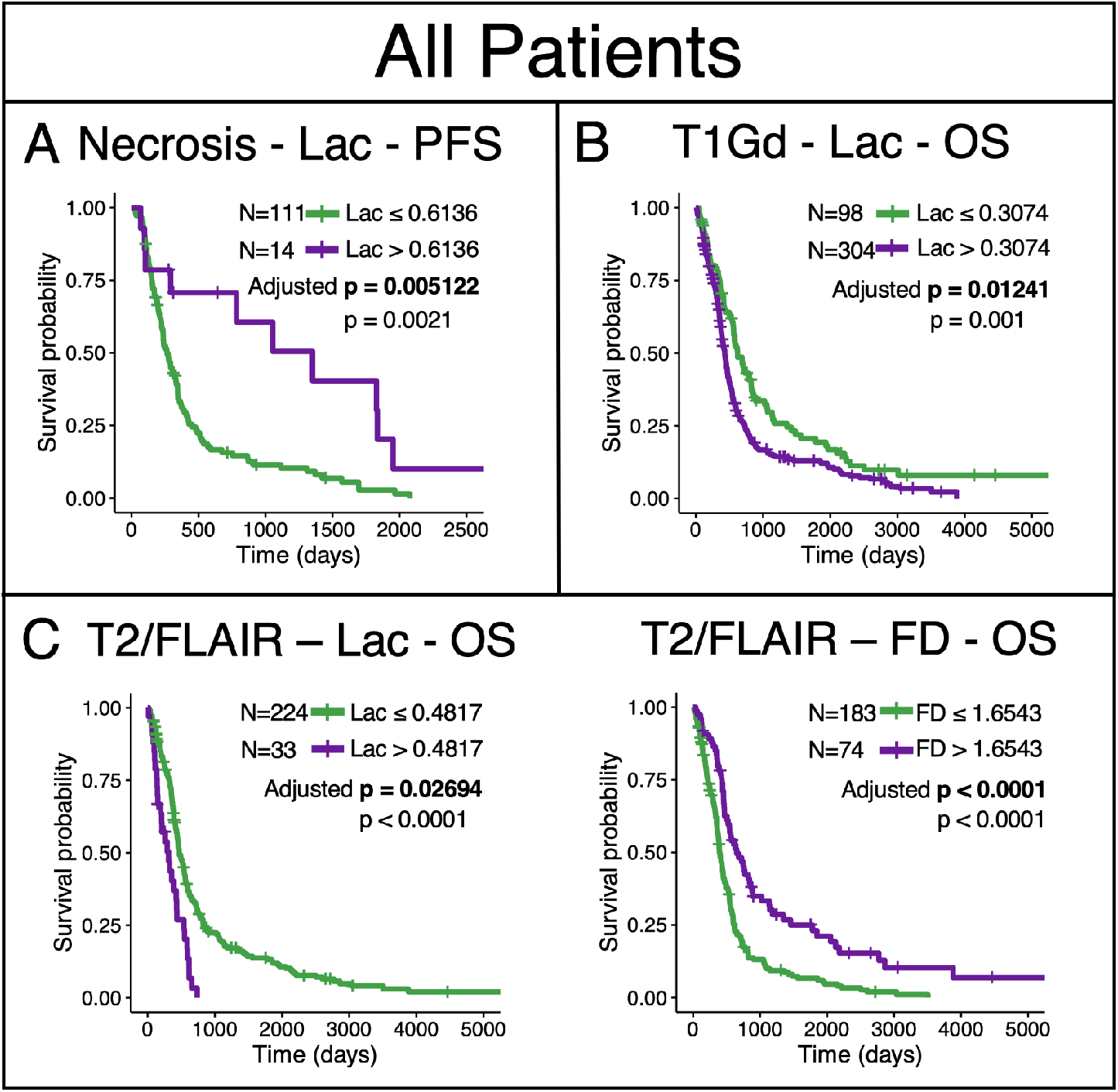
Results amongst all patients that remained significant with adjustment. **(A)** High lacunarity values of pretreatment necrosis lead to significantly longer PFS. A ‘ cutoff of 0.6136 maximally distinguished these groups. **(B)** Low lacunarity applied to T1Gd abnormalities was significantly better for OS, with 0.3074 chosen as the cutoff. **(C)** In T2/FLAIR abnormalities, lower lacunarity and higher FD values were associated with significantly improved overall survival.

#### T1Gd Region

In T1Gd regions, we observe a lacunarity threshold of 0.3074 that significantly distinguishes groups for overall survival with the group of lower lacunarity values surviving longer (adjusted p=0.012, n=402), see **Figure 2B**. We also saw that lacunarity distinguished PFS (unadjusted p=0.015) but this did not remain significant when adjusting for multiple comparisons (adjusted p=0.10).

#### T2/FLAIR Region

In T2/FLAIR abnormalities, lower lacunarity (Lac≤ 0.4817, adjusted p=0.0269) and higher FD values (FD>1.6543, adjusted p<0.0001) were associated with significantly improved overall survival (n=257), see **Figure 2C**.

### Current Standard of Care

We implemented the same analyses within a sub-cohort of patients known to have been treated with the current standard of care. Significant survival differences present in the whole cohort may not be reflected in this section due to a reduction in sample size leading to a reduction in statistical power.

#### Necrotic Region

In this subcohort of patients who received the current standard of care, we find that lacunarity and fractal dimension significantly distinguish overall survival and progression free survival. Lacunarity can distinguish progression free survival while adjusting for multiple comparisons (adjusted p=0.017, n=86), with the same threshold chosen to separate the groups as was chosen in the whole cohort (**Figure 2A** and **Figure 3**). We also see that the fractal dimension of necrosis can significantly distinguish overall survival (adjusted p=0.012, n=135) and progression free survival (adjusted p=0.018, n=86) (**Figure 3**) while accounting for multiple comparisons, with lower values conferring the survival benefit.

**Fig 3.**
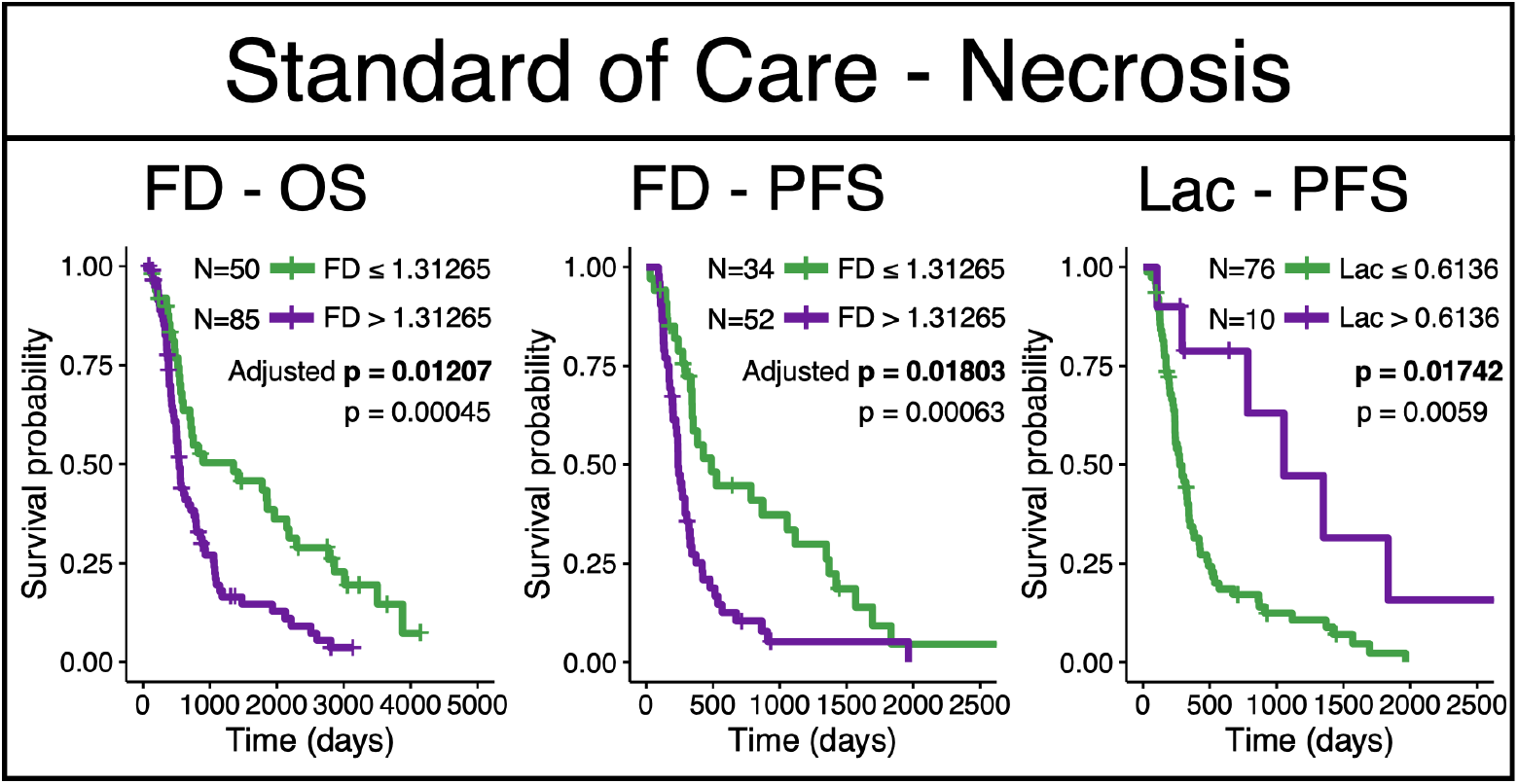
Fractal dimension and lacunarity cutoffs that significantly distinguish survival amongst patients confirmed to have received the current standard of care. Fractal dimension of necrosis regions significantly distinguished OS and PFS, while lacunarity significantly distinguished PFS. All three results remained significant with adjustment. Adjusted p values shown in bold.

#### T1Gd Region

Although no results held in this subcohort after adjusting for multiple comparisons, we did observe lacunarity thresholds that distinguished both overall and progression free survival with unadjusted significance (OS: p=0.043, adjusted p=0.31, PFS: p=0.016, adjusted p=0.091).

#### T2/FLAIR Region

Although no results held when adjusting for multiple comparisons in this subcohort, we did observe similar optimal cutoffs to those found in the larger cohort for both lacunarity and fractal dimension that significantly distinguished overall survival without adjustment (lacunarity p=0.019, FD p=0.020), see **Table 2**. We also note that the optimal cutoff for fractal dimension distinguished progression free survival but this result did not hold after adjustment for multiple comparisons (p=0.013).

**Table 2.** Median lacunarity and fractal dimension tests that showed at least one significant cutoff that distinguishes survival. We show both overall survival (OS) and progression free survival (PFS) with those that were significant (unadjusted p<0.05) in light gray. The results that remained significant while adjusting for multiple comparisons are shown in darker gray (adjusted p<0.05). The numerical value in the cell represents the optimal threshold of analyses that reached a level of significance.

|  |  | Necrosis |  | T1Gd |  | T2/FLAIR |  |
| --- | --- | --- | --- | --- | --- | --- | --- |
|  |  | Lac | FD | Lac | FD | Lac | FD |
| All Patients | OS | 0.27915 |  | 0.3074 |  | 0.4817 | 1.6543 |
|  | PFS | 0.6136 | 1.31265 | 0.3900 |  |  |  |
| Current SOC | OS | 0.28125 | 1.31265 | 0.3980 |  | 0.456 | 1.6621 |
|  | PFS | 0.6136 | 1.31265 | 0.3923 |  |  | 1.5202 |

We present a summary of the cutoffs that were found to be significant in either an unadjusted log-rank test or with adjustment (as described in Lausen and Schumacher (25)) in **Table 2**. We also present sex-specific breakdowns of these analyses in **Supplement 1**.

### Univariate and Multivariate Cox Proportional Hazard Models

We implemented univariate and multivariate Cox proportional hazard (CPH) models against overall survival and progression free survival. For overall survival, we present the univariate CPH models for the necrosis, T1Gd, and T2/FLAIR regions for age at diagnosis, fractal dimension, lacunarity, and tumor radius at presentation in **Figure 4**. Multivariate CPH models assessing the relationship between overall survival and lacunarity, age at diagnosis, and tumor radius are on the left side of **Figure 5** and multivariate analyses of fractal dimension, age at diagnosis and tumor radius are on the right side of **Figure 5**. We find that lacunarity and fractal dimension values of T2/FLAIR regions are the most commonly significant in univariate and multivariate analyses. The values found in **Figures 4–5** are presented in tables in **Supplement 3**. We chose to run two separate multivariate CPH models, one with fractal dimension and another with lacunarity, to test their independent ability as prognostic indicators against other factors. We present all the equivalent results amongst patients known to have received the current SOC in **Supplement 4**, which also show significance of both fractal dimension and lacunarity of T2/FLAIR for overall survival. We found that in a multivariate CPH analysis for progression-free survival of lacunarity of necrosis, necrosis radius, and age at diagnosis, that lacunarity and the radius were significant. No other variables were found to be significant for progression-free survival in either univariate or multivariate CPH analyses for any regions. We present plots of these results in **Supplement 5**.

**Fig 4.**
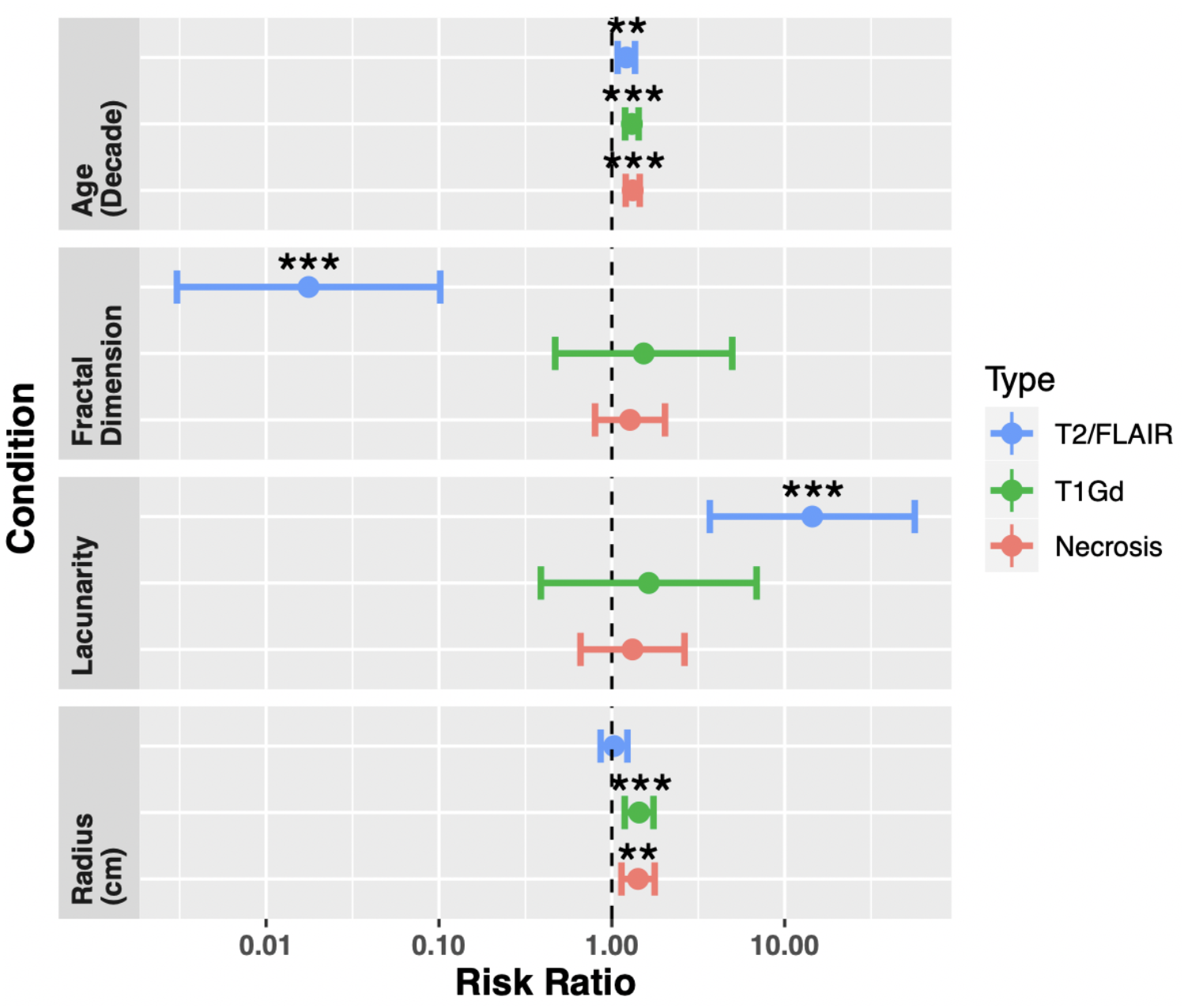
Univariate Cox proportional hazard models for age at diagnosis, fractal dimension, lacunarity and tumor radius, across all three regions (necrosis n=390, T1Gd n=402, T2/FLAIR n=257). Both fractal dimension and lacunarity of T2/FLAIR abnormalities are significant prognostic indicators of overall survival. Age at diagnosis was significant for all three regions, while radius was significant for both necrosis and T1Gd. Values of these tests can be found in **Supplement 3**.

**Fig 5.**
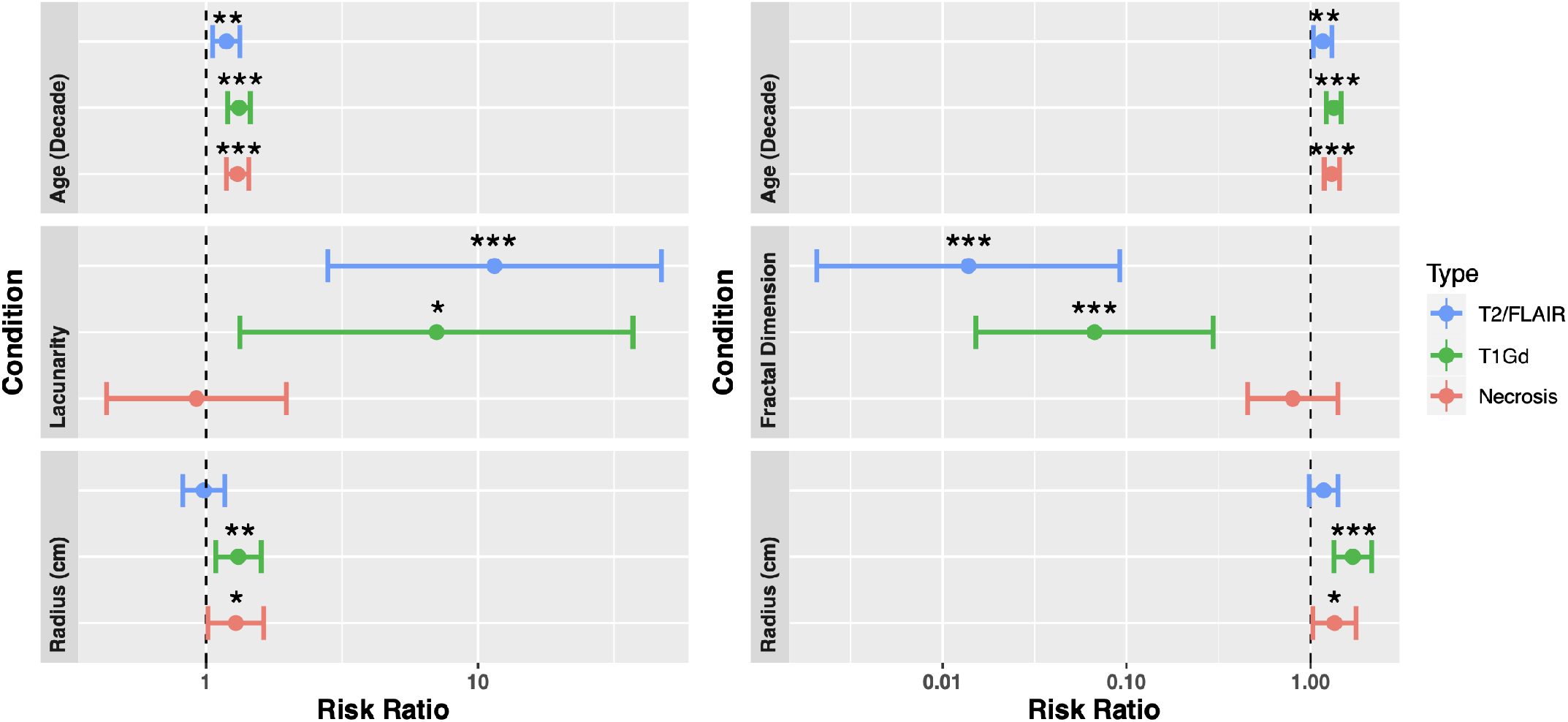
**(Left)** Multivariate CPH models of lacunarity, age at diagnosis, and abnormality radius for necrosis (N=390), T1Gd (N=402), and T2/FLAIR regions (N=257). Lacunarity of both T2/FLAIR and T1Gd were significant predictors of overall survival in their respective models (p=0.0007 and p=0.042, respectively). Age at diagnosis, as expected, was consistently significant for survival. Radius was a significant predictor for regions of necrosis and T1Gd (p=0.036). **(Right)** Corresponding Cox proportional hazard model of fractal dimension, age at diagnosis, and abnormality volumes for necrosis, T1Gd, and T2/FLAIR regions. Fractal dimension values of both T2/FLAIR and T1Gd regions were significant predictors of overall survival in their respective CPH models (p<0.0001 and p=0.0003, respectively). Age at diagnosis was consistently significant across all CPH models, while only T1Gd and necrosis radii were significant (p=0.0018 and p=0.028, respectively). For a complete table including confidence intervals and significance values, see **Supplement 3**.

### Correlations with other variables

We see significant negative correlations between lacunarity and fractal dimension in all of the imaging abnormalities tested (necrosis R=-0.55 p<0.0001, T1Gd R=-0.45, p<0.0001, T2/FLAIR R=-0.55, p<0.0001). With the exception of lacunarity of T1Gd regions (p=0.08), both lacunarity and fractal dimension are consistently significantly positively correlated with their corresponding volumes (all tests p<0.001).

Within the cohort of patients for which we have all three regions available (n=250), we observe significant positive correlations between both lacunarity and fractal dimension values of necrotic regions and T1Gd regions (both tests p<0.0001 Pearson). Significant positive correlations are also present between these metrics calculated on T1Gd regions and T2/FLAIR regions (lacunarity p=0.009 and FD p=0.018). We did not observe significant correlations between these metrics calculated on necrotic regions and T2/FLAIR abnormalities (lacunarity p=0.185 and FD p=0.339).

We see significantly lower fractal dimension values in necrosis-related abnormalities compared with their counterparts with enhancement on T1Gd (p<0.0001 t-test) and T2/FLAIR (p<0.001 t-test). No significant difference is observed in the fractal dimension between T1Gd and T2/FLAIR (p=0.49, t-test). We see significance between all three regions in lacunarity. Lacunarity is significantly higher in T2 than T1Gd enhancement (p<0.001, t-test), and necrosis is significantly higher than T2 enhancing regions (p<0.001, t-test). **Figure 6** shows boxplots of these values with their significant relationships highlighted.

**Figure 6:**
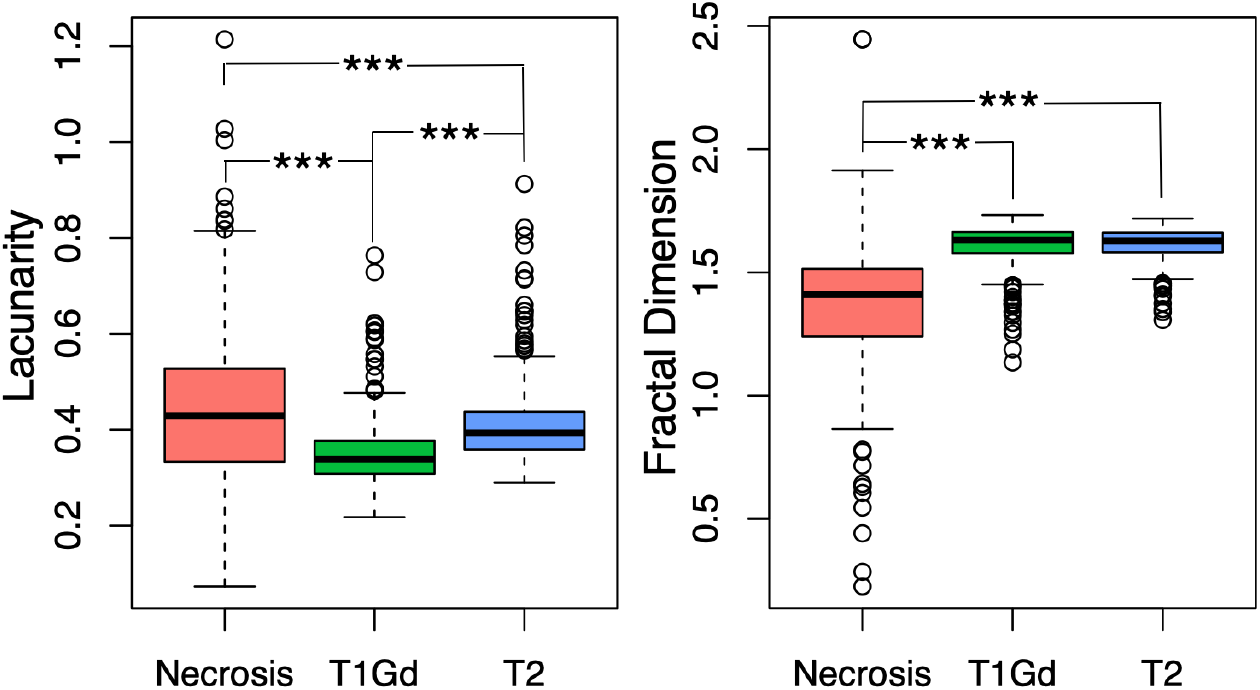
Significant differences in lacunarity and fractal dimension between regions of interest. **(Left)** We note significant differences in lacunarity values across all three regions of interest, with necrosis the highest, followed by T2/FLAIR, followed by T1Gd with necrosis (all comparisons p<0.001). **(Right)** FD values are significantly higher in T2/FLAIR and T1Gd when compared with necrosis (p<0.001) but we did not observe significant differences in FD between T2/FLAIR and T1Gd (p=0.49).

We note significant correlations of both morphological metrics with age at diagnosis. We observe a weak negative significant correlation between lacunarity and age at diagnosis in T1Gd enhancing regions (p=0.0217, R=-0.11 Pearson) but a positive significant correlation in T2/FLAIR enhancement (p=0.0035, R=0.18, Pearson). In contrast to this, we note a significant positive correlation between fractal dimension and age at diagnosis in T1Gd enhancing regions (p<0.001, R=0.19, Pearson) and a significant negative correlation in T2/FLAIR enhancing regions (p<0.001, R=-0.21, Pearson). We do not observe significant correlations within necrotic regions of lacunarity or fractal dimension with age at diagnosis. These results are shown in **Figure 7**.

**Figure 7:**
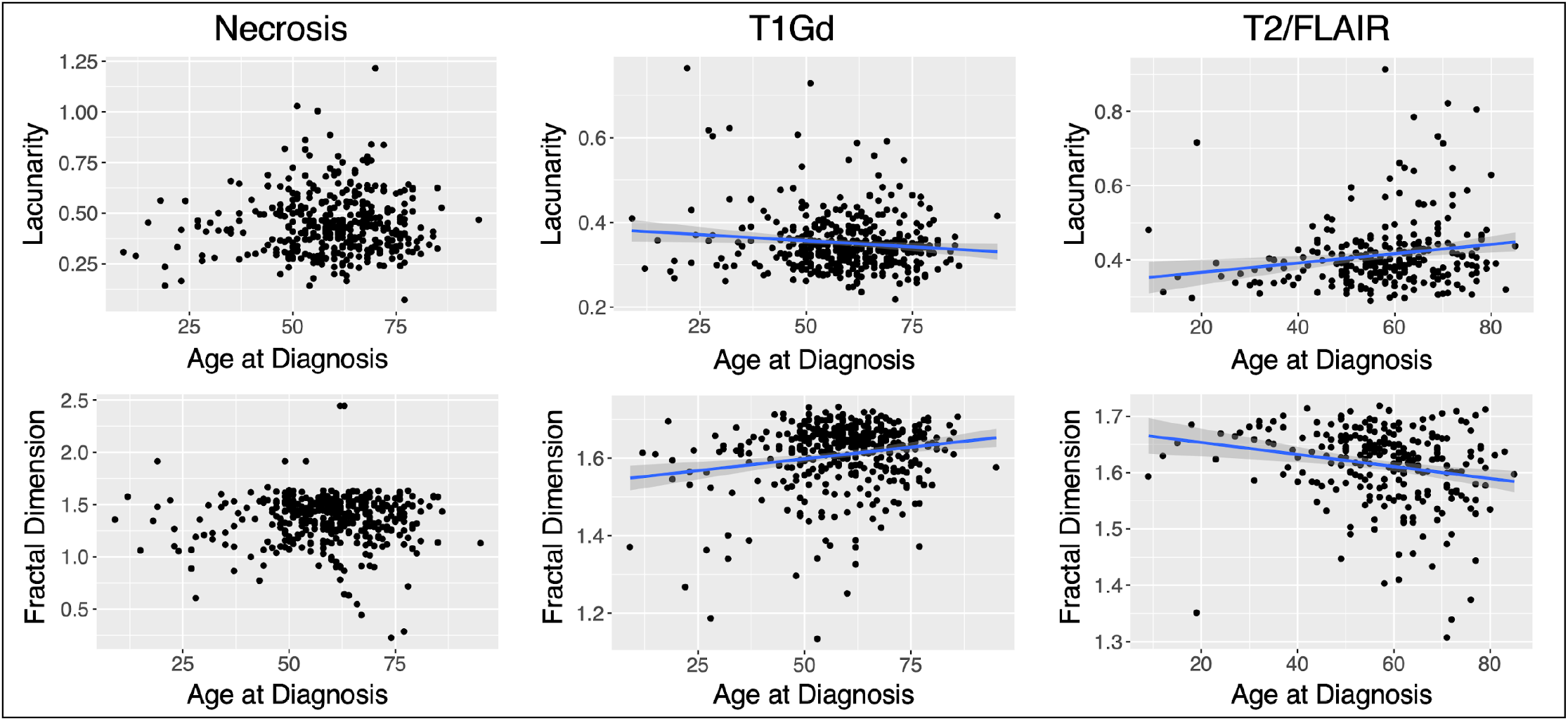
Significant relationships between morphology and age reversed between T1Gd and T2/FLAIR regions. **(Top row)** We see a weak significantly negative correlation between lacunarity and age at diagnosis in T1Gd ROIs (p=0.0217, R=-0.11) and a weak significantly positive correlation within T2/FLAIR ROIs (p=0.0035, R=0.18). **(Bottom row)** We see weak significant relationships between fractal dimension and age at diagnosis within T1Gd (p<0.001, R=0.19) and T2/FLAIR (p<0.001, R=-0.21) ROIs, respectively. Fractal dimension of T1Gd is positively correlated with age at diagnosis whereas the fractal dimension of T2/FLAIR ROIs has a negative correlation. Trend lines are shown for significant correlations.

## Discussion

Although clinical care teams will holistically consider multiple factors to determine the best course of action for each patient with GBM, they are somewhat limited in the computational tools and prognostic indicators that are readily available to them. The limited opportunities for tissue collection leads to a clinical reliance on imaging to make these decisions and an opportunity to maximize the utility of this information through prognostic imaging-derived biomarkers. Our results suggest that there is some prognostic impact of the shape of GBM at presentation. In contrast to the previous publication on this topic (16), we see some opposing relationships with these morphological metrics and patient survival for necrotic regions, in that our results generally show a low fractal dimension in necrotic regions is better for patient survival. To compare more closely with the previous publication, we also ran survival analyses for the mean lacunarity and fractal dimension values on necrotic regions. These metrics did not distinguish overall survival and progression free survival in as many instances as the median but still resulted in the same relationships where significant. We present a summary of these results in **Supplement 2**.

Within necrotic regions, we see more significant relationships between these morphological metrics and patient survival within patients who received the current standard of care. In this setting, FD can distinguish overall survival and progression free survival, and lacunarity can distinguish progression free survival while adjusting for multiple comparisons. Lacunarity also significantly distinguished overall survival but this result did not hold when adjusting for multiple comparisons.

The lacunarity of T2/FLAIR MRI pretreatment lesions has been shown to distinguish glioma grade (13). We have extended on this result to suggest that the shape of these regions also contain information on patient survival within GBM. Notably, in our cohort, we observe a benefit to OS of low T2/FLAIR lacunarity values and high T2/FLAIR fractal dimension values, both within our optimal threshold analysis and as continuous variables in univariate and multivariate CPH analyses. Within patients receiving the current standard of care, we also see that lacunarity and fractal dimension of T2/FLAIR abnormalities act as independent prognostic variables against age at diagnosis and the T2/FLAIR abnormality radius.

We note that although lacunarity and fractal dimension almost always significantly correlate with their associated volumes, they present different information that can be more significant for overall survival, particularly in T2/FLAIR regions. Rather unexpectedly, we also note that lacunarity and fractal dimension weakly correlate with patient age for T1Gd and T2/FLAIR regions. Machine learning has been used to reliably predict patient age from brain MRI of healthy adults (28), but to our knowledge no work has noted relationships between patient age and brain tumor size/shape.

It is important to note that a lack of statistical significance in survival analyses does not necessarily mean a lack of signal. In stratifying our patient cohorts, our statistical power to observe potential differences decreases. We chose to present optimal thresholds that did not remain significant while adjusting for multiple comparisons to show that we do see some signal with these morphological metrics in most cases. We hope that in the future these optimal thresholds will be tested in an independent patient cohort to validate the results presented here.

There has been some recent research showing that GBM location within the brain can impact outcome (26,27). Future work may explore lacunarity and fractal dimension against the location of the tumor, to determine whether location compliments or drives prognostic signals of lacunarity and fractal dimension that we have observed here. Further work may also explore the dynamics of these morphological markers throughout treatment and tumor progression.

Throughout this work, we have found relationships between the shape of segmented tumor regions and survival metrics. We found lacunarity and fractal dimension thresholds that significantly distinguish patient OS and PFS in our cohort, and showed that these act as continuous predictors of survival in some cases. These results warrant further investigation into the biological and genetic drivers behind the morphological presentation of GBM in a pretreatment setting.

## Author Contribution List

L.C. - Computed all morphological values, analyzed data, and wrote the manuscript. P.W. - Analyzed data and wrote the manuscript. H.W. - Data collection. M.M.M. - Provided clinical expertise. L.S.H. - Provided clinical expertise and supervised the project. K.R.S. - Supervised the project

## Abbreviations

GBM: Glioblastoma
FD: Fractal dimension
MRI: Magnetic resonance imaging
FLAIR: Fluid-attenuated inversion recovery
T1Gd: Gadolinium-enhanced T1-weighted
OS: Overall survival
PFS: Progression free survival

## Acknowledgements

The authors gratefully acknowledge the funding that made this research possible from the NIH (R01NS060752, R01CA164371, U54CA143970, U54CA193489, U01CA220378, U54CA210180), the Arizona Biomedical Research Commission (ADHS16-162514), the James S. McDonnell Foundation, and the Ben and Catherine Ivy Foundation. The authors also acknowledge the image analysis team for their work segmenting tumors.

