## Supplementary Material for "Morphological Metrics of Magnetic Resonance Imaging of Glioblastoma as Biomarkers of Prognosis"

**Co-senior authors contributed equally

**Author Contribution List:** L.C. - Computed all morphological values, analyzed data, and wrote the manuscript. P.W. - Analyzed data and wrote the manuscript. H.W. - Data collection. M.M.M. - Provided clinical expertise. L.S.H. - Provided clinical expertise and supervised the project. K.R.S. - Supervised the project

**Running Title:** Morphological Metrics as Pretreatment Biomarkers in Glioblastoma

**Keywords:** Lacunarity, Fractal Dimension, Glioblastoma, Survival, Morphology

**Abbreviations:** Glioblastoma (GBM). Fractal dimension (FD). Magnetic resonance imaging (MRI). Fluid-attenuated inversion recovery (FLAIR). Gadolinium-enhanced T1-weighted (T1Gd). Overall survival (OS). Progression free survival (PFS).

**Abstract:**
Morphological characteristics have been linked to outcomes across a variety of cancers. Lacunarity is a quantitative morphological measure of how shapes fill space while fractal dimension is a morphological measure of the complexity of pixel arrangement. Glioblastoma is the most aggressive primary brain tumor with a short expected survival given the current standard-of-care treatment. Due to the sensitive location of the tumor, there is a heavy reliance on imaging to assess the state of the disease in the clinic. In this project, we computed lacunarity and fractal dimension values for glioblastoma-induced abnormalities on gadolinium-enhanced T1-weighted magnetic resonance imaging (T1Gd MRI) as well as T2-weighted (T2) and fluid-attenuated inversion recovery (FLAIR) MRIs. In our patient cohort (n=402), we aim to connect these morphological metrics calculated on pretreatment MRI with the survival of patients with GBM. We calculated lacunarity and fractal dimension across all MRI slices on necrotic regions (n=390) and abnormalities on T1Gd MRI (n=402), as well as on enhancing abnormalities present on T2/FLAIR MRI (n=257). We also explored the relationship between these metrics and age at diagnosis, as well as abnormality volume. We found statistically significant relationships to outcome across all three imaging subtypes, with the shape of T2/FLAIR abnormalities showing the strongest relationship with overall survival. The link between morphological and survival metrics could be driven by underlying biological phenomena, tumor location or microenvironmental factors that should be further explored.

### Supplemental Data

**Supplement 1: Sex-specific Results**

All results presented in the previous subsections were carried out on a sex-specific basis.

| | | Necrosis | | | T1Gd Enhancing | | | T2/FLAIR | | |
|---|---|---|---|---|---|---|---|---|---|---|
| | | All | Male | Female | All | Male | Female | All | Male | Female |
| Everyone | OS | 390 | 246 | 144 | 402 | 253 | 149 | 257 | 164 | 93 |
| | PFS | 125 | 84 | 41 | 130 | 87 | 43 | 78 | 50 | 28 |
| Current SOC | OS | 135 | 88 | 47 | 142 | 92 | 50 | 92 | 56 | 36 |
| | PFS | 86 | 58 | 28 | 89 | 59 | 30 | 56 | 33 | 23 |

**Cohort numbers for patients with known overall survival (OS) and progression free survival (PFS).** This table shows the patients with known OS and PFS, including the subsets known to have received the current standard of care (SOC). We also present how these cohorts are split by patient sex. The discrepancy between patients with necrosis ROIs and T1Gd enhancing is due to 10 patients with negligible necrosis that did not meet our criteria to be included in this retrospective study.

| | | Necrosis | | | T1Gd Enhancing | | | T2/FLAIR | | |
|---|---|---|---|---|---|---|---|---|---|---|
| | | All | Male | Female | All | Male | Female | All | Male | Female |
| Everyone | OS | 0.27915 | 0.33525 | 0.2919 | 0.3074 | 0.3074 | 0.30305 | 0.4817 | 0.4817 | 0.4705 |
| | PFS | 0.6136 | 0.6136 | | 0.39 | | 0.3893 | | | |
| Current SOC | OS | 0.28125 | | | 0.398 | | | 0.456 | | 0.37355 |
| | PFS | 0.6136 | | 0.6011 | 0.3923 | | 0.3816 | 0.34695 | | |

**Median lacunarity tests that showed at least one significant cutoff that distinguishes survival.** We show both overall survival (OS) and progression free survival (PFS) with those that were significant in light gray. The results that remained significant while adjusting for multiple comparisons are shown in dark gray.

| | | Necrosis | | | T1Gd Enhancing | | | T2/FLAIR | | |
|---|---|---|---|---|---|---|---|---|---|---|
| | | All | Male | Female | All | Male | Female | All | Male | Female |
| Everyone | OS | | | 1.26725 | | | 1.565 | 1.6543 | 1.64625 | 1.6541 |
| | PFS | 1.31265 | 1.31265 | | | | | 1.62515 | | 1.62305 |
| Current SOC | OS | 1.31265 | 1.31265 | 1.26725 | | 1.675 | 1.565 | 1.6621 | 1.51575 | 1.66355 |
| | PFS | 1.31265 | 1.31265 | | | | | 1.62515 | | |

**Median fractal dimension tests that showed at least one significant cutoff that distinguishes survival.** We show both overall survival (OS) and progression free survival (PFS) with those that were significant in light gray. The results that remained significant while adjusting for multiple comparisons are shown in dark gray.

#### Supplement 2: Testing Mean Lacunarity and Mean Fractal Dimension

| | | Necrosis | | | T1Gd Enhancing | | | T2/FLAIR | | |
|---|---|---|---|---|---|---|---|---|---|---|
| | | All | Male | Female | All | Male | Female | All | Male | Female |
| Everyone | OS | 0.3405833 | 0.3405833 | | 0.3051281 | 0.3039867 | 0.3016833 | 0.4915923 | 0.4504357 | 0.4820333 |
| | PFS | 0.606835 | | | | | | 0.3495091 | 0.3495091 | 0.3699375 |
| Current SOC | OS | | | | 0.3069367 | 0.3069367 | | 0.4820333 | 0.4915923 | |
| | PFS | 0.6120233 | | | | 0.2833917 | 0.3824581 | 0.3540625 | 0.3540625 | |

**Mean lacunarity tests that showed at least one significant cutoff that distinguishes survival.** We show both overall survival (OS) and progression free survival (PFS) with those that were significant in light gray. The results that remained significant while adjusting for multiple comparisons are shown in dark gray.

| | | Necrosis | | | T1Gd Enhancing | | | T2/FLAIR | | |
|---|---|---|---|---|---|---|---|---|---|---|
| | | All | Male | Female | All | Male | Female | All | Male | Female |
| Everyone | OS | | | 1.348725 | | | 1.547462 | 1.624938 | 1.624938 | |
| | PFS | 1.251472 | 1.238051 | | | | | 1.622186 | | |
| Current SOC | OS | 1.2525 | | | | | 1.5958 | 1.650638 | | 1.6441 |
| | PFS | 1.251472 | 1.238051 | 1.251472 | | | 1.587316 | 1.608108 | | |

**Mean fractal dimension tests that showed at least one significant cutoff that distinguishes survival.** We show both overall survival (OS) and progression free survival (PFS) with those that were significant in light gray. The results that remained significant while adjusting for multiple comparisons are shown in dark gray.

#### Supplement 3: Values Presented in Cox Proportional Hazard Plots

| **Necrosis Variables (p value, HR, 95% CI)** | |
|---|---|
| **Univariate CPH Outcomes** | |
| Lacunarity | 0.435, 1.31732822, [0.659038963 2.6331579] |
| Fractal Dimension | 0.307, 1.27443832, [0.799949159 2.0303703] |
| Age (decades) | **<0.0001, 1.32074821, [1.203296409 1.4496643]** |
| Radius (cm) | **0.00194, 1.41831613, [1.137130169 1.7690329]** |
| **Multivariate CPH Outcomes** | |
| Lacunarity | 0.8351, 0.9222878, [0.4305380 1.975702] |
| Age (decades) | **<0.0001, 1.3048678, [1.1867259 1.434771]** |
| Radius (cm) | **0.0364, 1.2872644, [1.0161378 1.630733]** |

| | |
|---|---|
| Fractal Dimension | 0.441, 0.80070990, [0.454778154 1.40977825] |
| Age (decades) | **<0.0001, 1.30563149, [1.187578119 1.43542017]** |
| Radius (cm) | **0.028, 1.35074734, [1.032977624 1.76627096]** |

| **T1Gd Variables (p value, HR, 95% CI)** | |
|---|---|
| **Univariate CPH Outcomes** | |
| Lacunarity | 0.503, 1.63494814, [0.388452428 6.8812942] |
| Fractal Dimension | 0.48, 1.52922637, [0.470292430 4.9725089] |
| Age (decades) | **<0.0001, 1.30594705, [1.193211710 1.4293337]** |
| Radius (cm) | **0.0002, 1.43847096, [1.187303246 1.7427719]** |
| **Multivariate CPH Outcomes** | |
| Lacunarity | **0.02143, 7.0339701, [1.3345471 37.073803]** |
| Age (decades) | **<0.0001, 1.3229636, [1.2037698 1.453960]** |
| Radius (cm) | **0.00552, 1.3147489, [1.0836820 1.595085]** |
| Fractal Dimension | **0.000356, 0.06702770, [0.015206290 0.29545094]** |
| Age (decades) | **<0.0001, 1.33902267, [1.219116419 1.47072230]** |
| Radius (cm) | **<0.0001, 1.69721089, [1.338093669 2.15270789]** |

| **T2/FLAIR Variables (p value, HR, 95% CI)** |
|---|

| **Univariate CPH Outcomes** | |
|---|---|
| Lacunarity | **0.00013, 14.44702788, [3.689517402 56.5701668]** |
| Fractal Dimension | **<0.0001, 0.01760181, [0.003048264 0.1016394]** |
| Age (decades) | **0.0011, 1.21317106, [1.080767757 1.3617949]** |
| Radius (cm) | 0.746, 1.02989506, [0.861678973 1.2309501] |
| **Multivariate CPH Outcomes** | |
| Lacunarity | **0.000693, 11.4824765, [2.8026852 47.043195]** |
| Age (decades) | **0.003493, 1.1885148, [1.0584508 1.334561]** |
| Radius (cm) | 0.844741, 0.9824919, [0.8232941 1.172473] |
| Fractal Dimension | **<0.0001, 0.01383143 0.002077994 0.09206398** |
| Age (decades) | **0.00873, 1.16620359 1.039598702 1.30822674** |
| Radius (cm) | 0.07375, 1.17864640 0.984331381 1.41132077 |

### Supplement 4: Current Standard of Care Cox Proportional Hazard Plots

#### Current SOC - Univariate Analysis - Overall Survival

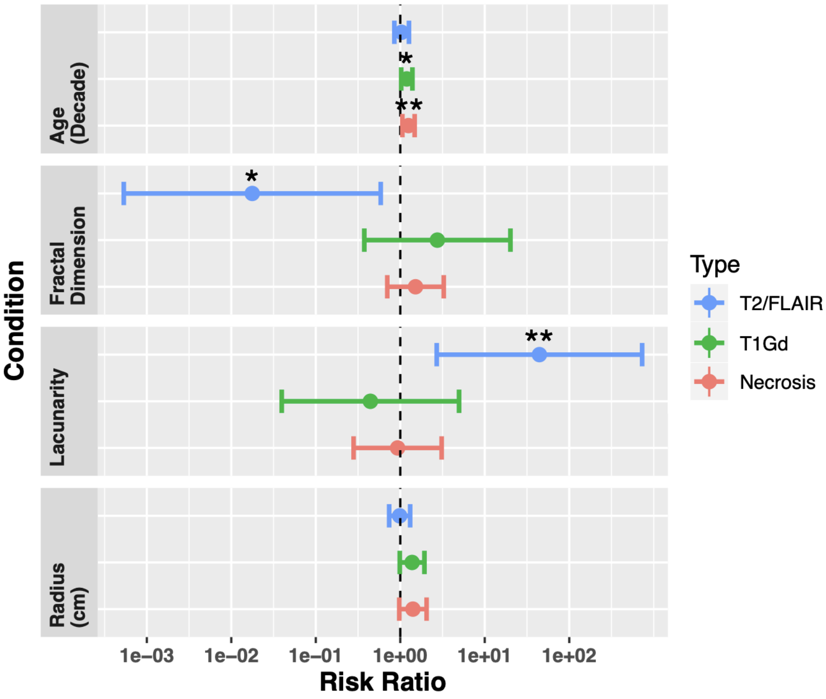


Univariate Cox proportional hazard model for overall survival of patients known to have received the current SOC. Patient counts are 135, 142, 92 for necrosis, T1Gd and T2/FLAIR analyses, respectively. Both fractal dimension and lacunarity are significant for T2/FLAIR regions. Age at diagnosis was significant only for necrosis and T1Gd regions. The lack of signal for age at diagnosis for T2/FLAIR could be owing to the lower sample size.

#### Current SOC - Multivariate Analysis - Overall Survival

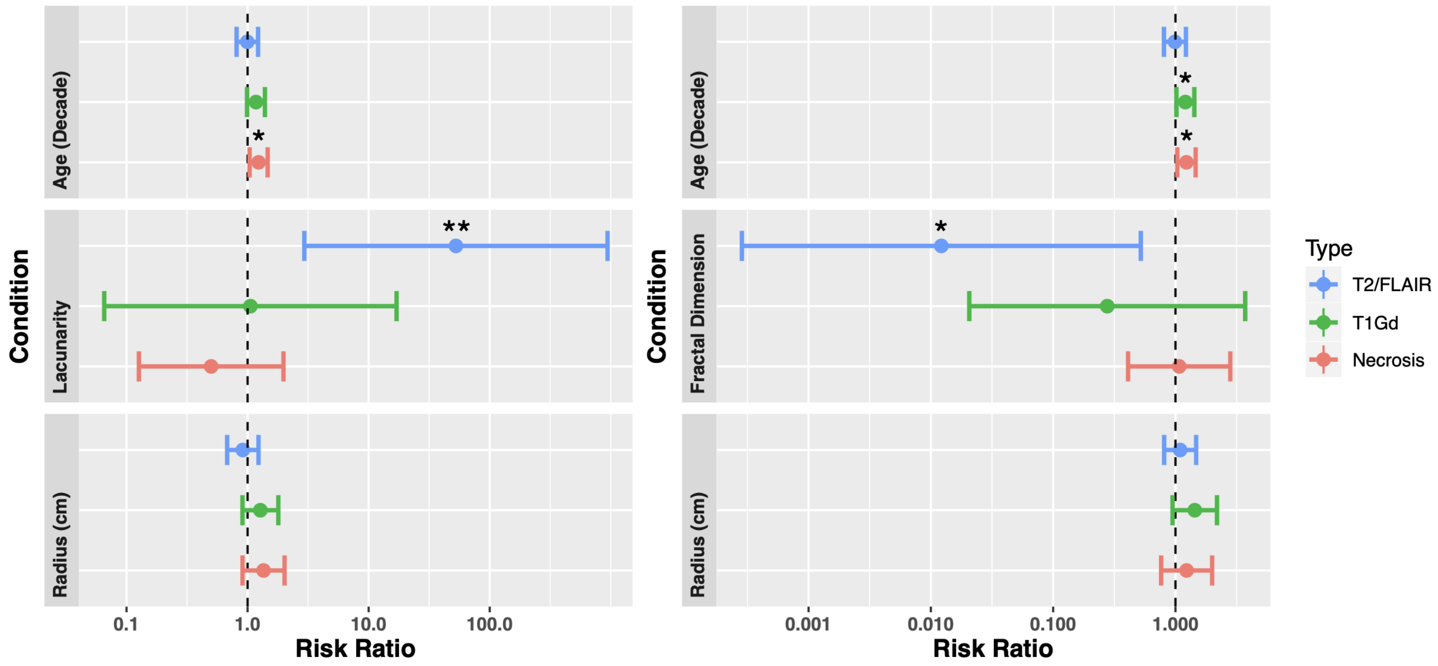


Multivariate Cox proportional hazard model for overall survival of patients known to have received the current standard of care. Separate analyses for necrosis (n=135), T1Gd (n=142) and T2/FLAIR (n=92) are all presented here. **(Left)** Lacunarity of T2/FLAIR regions showed significant influence for overall survival. The only other variable to show significance was age at diagnosis within the necrosis analysis. **(Right)** Fractal dimension was significant only for T2/FLAIR regions, with age at diagnosis significant in necrosis and T1Gd analyses.

#### Supplement 5: Progression-Free-Survival Cox Proportional Hazard Plots

##### Univariate CPH for Progression-Free Survival

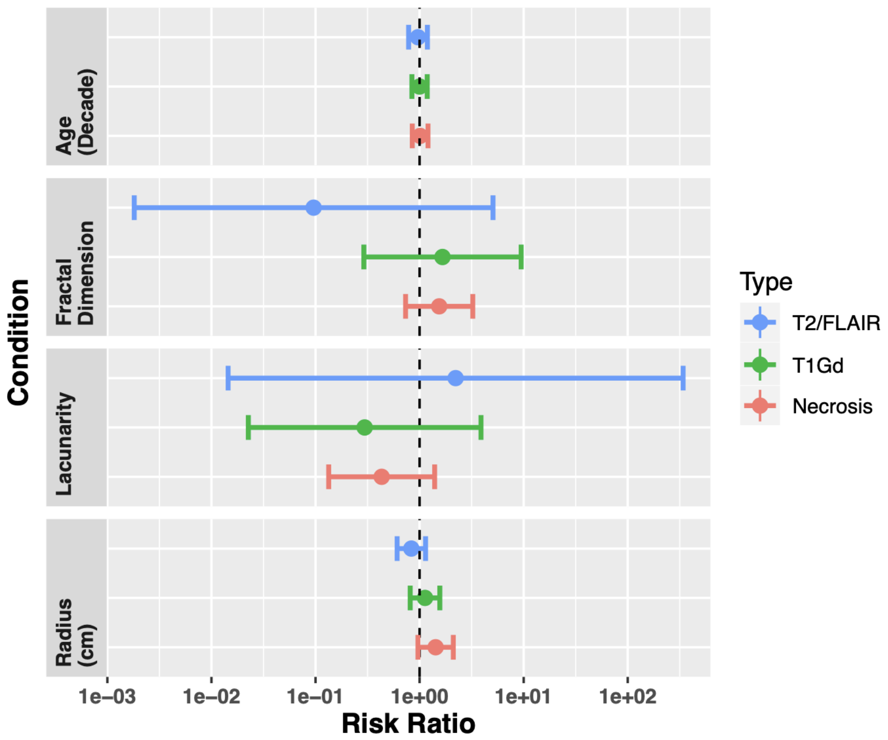


Univariate Cox proportional hazard model for progression-free survival of all patients. Patient counts are 125, 130 and 78 for necrosis, T1Gd and T2/FLAIR analyses, respectively. No variables showed significance in any of these analyses.

##### Multivariate CPH for Progression-Free Survival

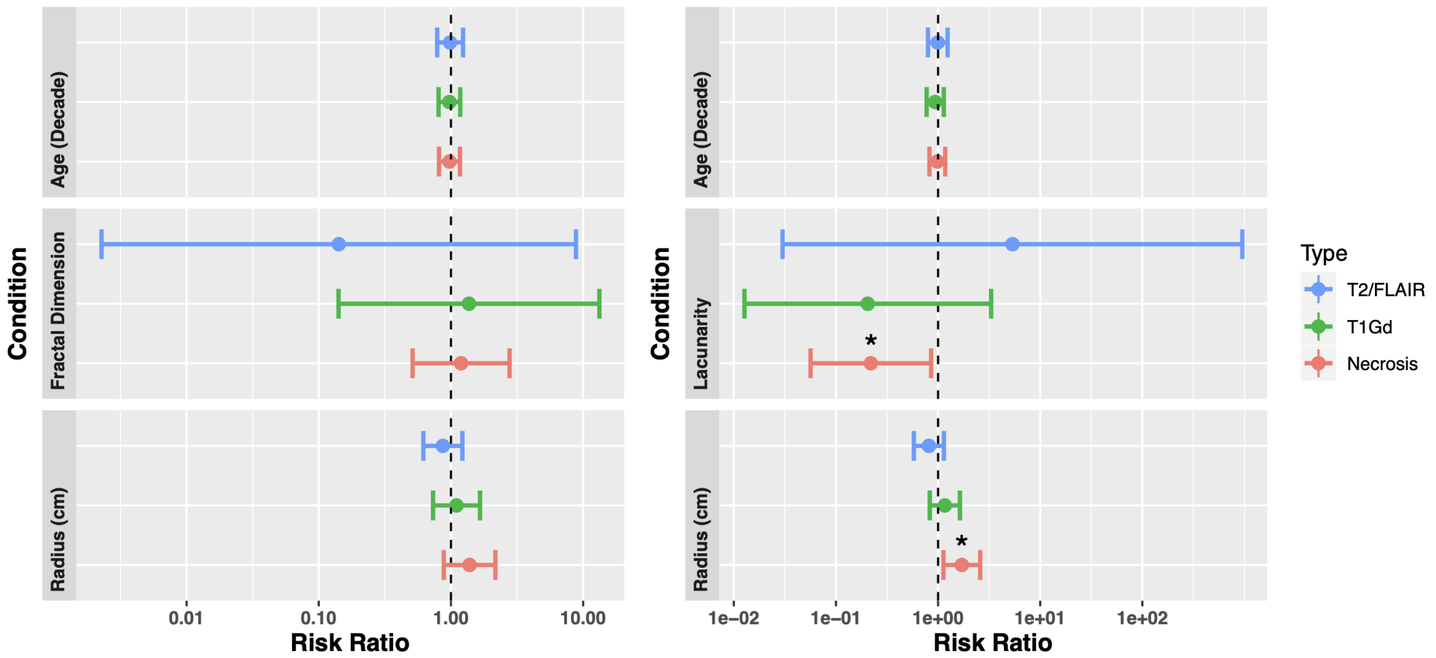


Multivariate Cox proportional hazard model for progression-free survival. Separate analyses for necrosis (n=125), T1Gd (n=130) and T2/FLAIR (n=78) are all presented here. **(Left)** No significance was found for any variables. **(Right)** Lacunarity of T1Gd regions showed significant influence for overall survival. The only other variable to show significance was necrosis radius.
